# Centrally generated movement-related signals engage ipsilateral S1 finger representations after tetraplegia

**DOI:** 10.64898/2026.09.03.748249

**Authors:** Finn Rabe, Hunter Schone, Paige Howell, Dario Pfyffer, Patrick Freund, Nicole Wenderoth, Sanne Kikkert

## Abstract

Ipsilateral primary somatosensory cortex (S1) exhibits somatotopically organized activity during unilateral hand movements, but whether this activity depends on peripheral input remains debated. We addressed this question using functional magnetic resonance imaging (fMRI) in 14 individuals with tetraplegia and 18 able-bodied controls. Participants performed or attempted individual finger movements during 3T fMRI. We quantified movement-related univariate activity and multivariate representational structure within ipsilateral S1. Univariate activity within the anatomically defined ipsilateral S1 hand area did not differ significantly between groups, whereas a continuous spatial analysis identified greater movement-related activity in individuals with tetraplegia in S1 bands lateral to the hand representation. Multivariate analyses revealed reliable separation of finger-specific activity patterns in both groups, with no significant group difference in mean inter-finger distance. The typicality of the overall representational geometry also did not differ significantly between groups. Finally, a person with complete hand paralysis and no measurable feeling or movement still showed clear, finger-specific brain activity patterns and a typical representation of the affected hand. Together, these findings indicate that centrally generated movement-related signals can engage finger-specific representations in ipsilateral S1 without overt movement or measurable hand-related peripheral input. Ipsilateral somatotopic activity therefore cannot be explained solely by sensory reafference from the moving hand.

## INTRODUCTION

During unilateral hand movements, S1 activity peaks in the hemisphere contralateral to the moving hand. However, a smaller yet reliable response is also observed in ipsilateral S1^1–5^. This ipsilateral S1 activity is somatotopically organized and emerges in the cortical territory that, in the opposite (contralateral) hemisphere, normally represents the same body part, the so-called homologous representation^1,6^. Importantly, although this S1 representation can be activated by movements of either hand, the neuronal populations engaged by ipsilateral hand movements are largely distinct from those activated during contralateral movements^7,8^.

The precise origins of ipsilateral S1 activation remain debated. Proposed sources include (1) bilateral sensorimotor body part representations that support complex movements^5,9^, (2) reduced interhemispheric inhibition of transcallosal pathways during unilateral movements^6,10–14^, (3) top-down inputs from higher order motor areas with bilateral organization^15^, (4) bilateral thalamocortical projections^16^, or (5) uncrossed peripheral afferents conveying sensory inputs directly to the ipsilateral hemisphere^17^. Dissecting the relative contributions of central versus peripheral sources is essential for determining how ipsilateral S1 activity arises during unilateral limb movements.

Several observations support a central contribution. For example, studies in both humans and non-human primates have demonstrated that ipsilateral activity is dynamically modulated by task context^5,18^, with ipsilateral representations emerging selectively under specific cognitive demands^19–21^. Furthermore, finger-specific patterns are more distinct during active than passive finger movements even when peripheral sensory signals reach contralateral S1 in both conditions^1,22^. Nevertheless, peripheral input was present in these studies, meaning that its contribution could not be excluded. Nevertheless, the relative reduction of ipsilateral finger information during passive stimulation indicates that ascending sensory input alone is insufficient to account for these representations and implicates central processes associated with active movement in their generation^1^.

While these studies consistently point to top-down, movement-related processes, as a major driver of ipsilateral S1 activity, hand-related peripheral input was always present, preventing its contribution from being ruled out. Active movement inherently combines centrally generated motor signals with the movement-related sensory feedback, meaning that their respective contributions cannot be fully separated in the intact nervous system^23^. Establishing whether central signals alone are sufficient to activate ipsilateral S1 representations therefore requires a clinical model in which motor signals can be generated centrally without overt movement or the associated hand-related sensory feedback.

Spinal cord injury (SCI) provides such a human model in which ascending sensory feedback from and motor function of body parts below the lesion level is absent or markedly reduced while the cortical itself remains intact ^24–28^. We, and others, previously showed that attempted finger movements can engage contralateral S1 finger representations in individuals without measurable hand function or detectable spared tissue bridges across the spinal lesion^29–33^. These preserved finger representations, which can be observed in the absence of peripheral inputs, must be driven by top-down mechanisms. Evidence concerning ipsilateral S1 is more limited. Recent intracortical recordings from an individual with incomplete tetraplegia, hence, not fully deafferented, showed that finger movements elicited high-gamma bursts in the ipsilateral S1 hand area^34^. Interestingly, this ipsilateral S1 activity emerged faster than would be expected if it were driven solely by sensory feedback travelling from the moving hand to the cortex. It was therefore argued that these ipsilateral responses likely reflect central, top-down signals rather than uncrossed peripheral afferents^34^. However, because peripheral pathways were not completely disrupted, this finding could not establish whether central signals alone are sufficient to elicit ipsilateral S1 representations. Moreover, the evidence was obtained from a single individual and did not establish whether the ipsilateral activity contained a typical finger-specific representational organization.

To tease apart peripheral and central contributions to ipsilateral S1 activity, we scanned tetraplegic participants using 3 tesla fMRI to investigate ipsilateral finger representations during visually cued unimanual individual finger movements. If an SCI individual was unable to perform overt finger movements, we asked them to attempt to perform the finger movement ^33^). Crucially, one of the tetraplegic participants had a complete SCI (i.e., no intact peripheral hand sensory inputs), allowing us to more finely test whether peripheral inputs are necessary to elicit ipsilateral finger-specific somatotopic activity in S1.

## RESULTS

We first examined the magnitude and spatial distribution of movement-related activity in bilateral S1 using univariate analyses, before assessing finger-pattern separability and the typicality of the representational geometry in ipsilateral S1 using multivariate analyses.

We investigated functional activity during unimanual (attempted) finger movements. Such movements involve the active generation of a motor command from the brain to the periphery, even if the signal does not result in muscle contraction due to the SCI^35^. Results pertaining to contralateral S1 presented here have been published previously^33,36^ and are included to facilitate interpretation and direct comparison with the new results taken from ipsilateral S1.

The study included 14 individuals with chronic tetraplegia with a diverse clinical profile (see Table 1). SCI completeness ranged from AIS-A to AIS-D, with neurological levels of injury spanning C2 to C7. Time since injury varied from 6 months to 33 years. Sensorimotor impairments were similarly heterogeneous, with bilateral GRASSP scores ranging between 21 and 220 (compared to a healthy benchmark score of 232). For comparison, we also assessed a control group of 18 healthy participants matched for age, sex, and handedness.

**Table 1.** Demographic and clinical characteristics of the tetraplegic participants. Remaining sensorimotor function was assessed using the Graded Redefined Assessment of Strength, Sensibility and Prehension (GRASSP), and participants are ordered by overall GRASSP score. The table additionally reports remaining midsagittal tissue width at the lesion level. These measurements were unavailable for two patients due to metal artifacts. Sex: F, female; M, male; age, years; AIS grade, American Spinal Injury Association Impairment Scale grade according to the International Standards for Neurological Classification of Spinal Cord Injury (ISNCSCI), where A = complete, B = sensory incomplete, C = motor incomplete, D = motor incomplete, and E = normal; neurological level of injury, according to the ISNCSCI; dominant hand, according to the Edinburgh Handedness Inventory, L = left and R = right; GRASSP maximum score, 232; tested side, side with the lower GRASSP score; motor and sensory GRASSP scores are shown for both hands. The total score includes prehension sub-tests not shown here. Adapted from ^58^.

| ID | Sex | Age | Years since injury | AIS grade | Cause of injury | Neurological level of injury | Dominant hand | GRASSP score | Hand tested | GRASSP tested Hand. motor/sensory | GRASSP other Hand. motor/sensory | Midsagittal tissue bridges width |
| --- | --- | --- | --- | --- | --- | --- | --- | --- | --- | --- | --- | --- |
| S01 | M | 32 | 4 | A | trauma | C4 | L | 21 | L | 9/0 | 8/4 | 0 |
| S02 | M | 52 | 32 | A | trauma | C5 | R | 78 | L | 16/5 | 13/16 | - |
| S03 | M | 35 | 4 | A | trauma | C5 | R | 90 | L | 16/17 | 16/13 | 0 |
| S04 | M | 41 | 19 | A | trauma | C5 | L | 105 | L | 19/11 | 20/22 | 0 |
| S05 | M | 52 | 10 | A | trauma | C3 | L | 118 | R | 23/0 | 29/0 | 2.08 |
| S06 | M | 67 | 26 | A | trauma | C7 | R | 119 | L | 29/2 | 33/4 | - |
| S07 | M | 57 | 33 | C | trauma | C6 | L | 145 | R | 25/19 | 23/22 | 1.28 |
| S08 | M | 67 | 4 | D | trauma | C5 | L | 173 | L | 41/9 | 32/15 | 1.97 |
| S09 | M | 59 | 12 | D | trauma | C4 | R | 187 | R | 43/9 | 45/15 | 2.28 |
| S10 | M | 42 | 2 | D | trauma | C3 | R | 187 | R | 41/13 | 42/15 | 2.22 |
| S11 | M | 58 | 0.5 | D | ischemic | C4 | R | 194 | R | 37/17 | 13/23 | 1.7 |
| S12 | F | 71 | 16 | D | trauma | C6 | R | 196 | R | 32/24 | 50/24 | 3.6 |
| S13 | M | 65 | 1 | D | trauma | C2 | R | 218 | R | 42/24 | 44/24 | 5.11 |
| S14 | M | 74 | 6 | D | surgery | C3 | L | 220 | R | 47/24 | 46/24 | 0.85 |

### MOVEMENT-RELATED ACTIVITY IS ELEVATED LATERAL TO THE IPSILATERAL S1 HAND AREA AFTER TETRAPLEGIA

First, we examined group-level activity maps per group. During unimanual finger tapping, we found that both the control and tetraplegic participants exhibited significant movement-related activity across sensorimotor areas, including in the sensorimotor cortex ipsi- and contralateral to the moved fingers (**Appx. A1**).

Next, to characterize the spatial distribution of movement-related activity across S1, we divided S1 into 50 bands and extracted the averaged fingers movement-related activity level per band (**Fig. 1**). In contralateral S1 (**Fig. 1A**), tetraplegic individuals and controls exhibited qualitatively similar activation level profiles, with strongest activity in the anatomically defined hand area (gray shaded region; individuals with tetraplegia: z = 5.8 ± 0.7; controls: z = 5.6 ± 0.8; p = 0.74) and no significant group differences in activity levels across bands (all bands z < 3.24).

**Figure 1.**
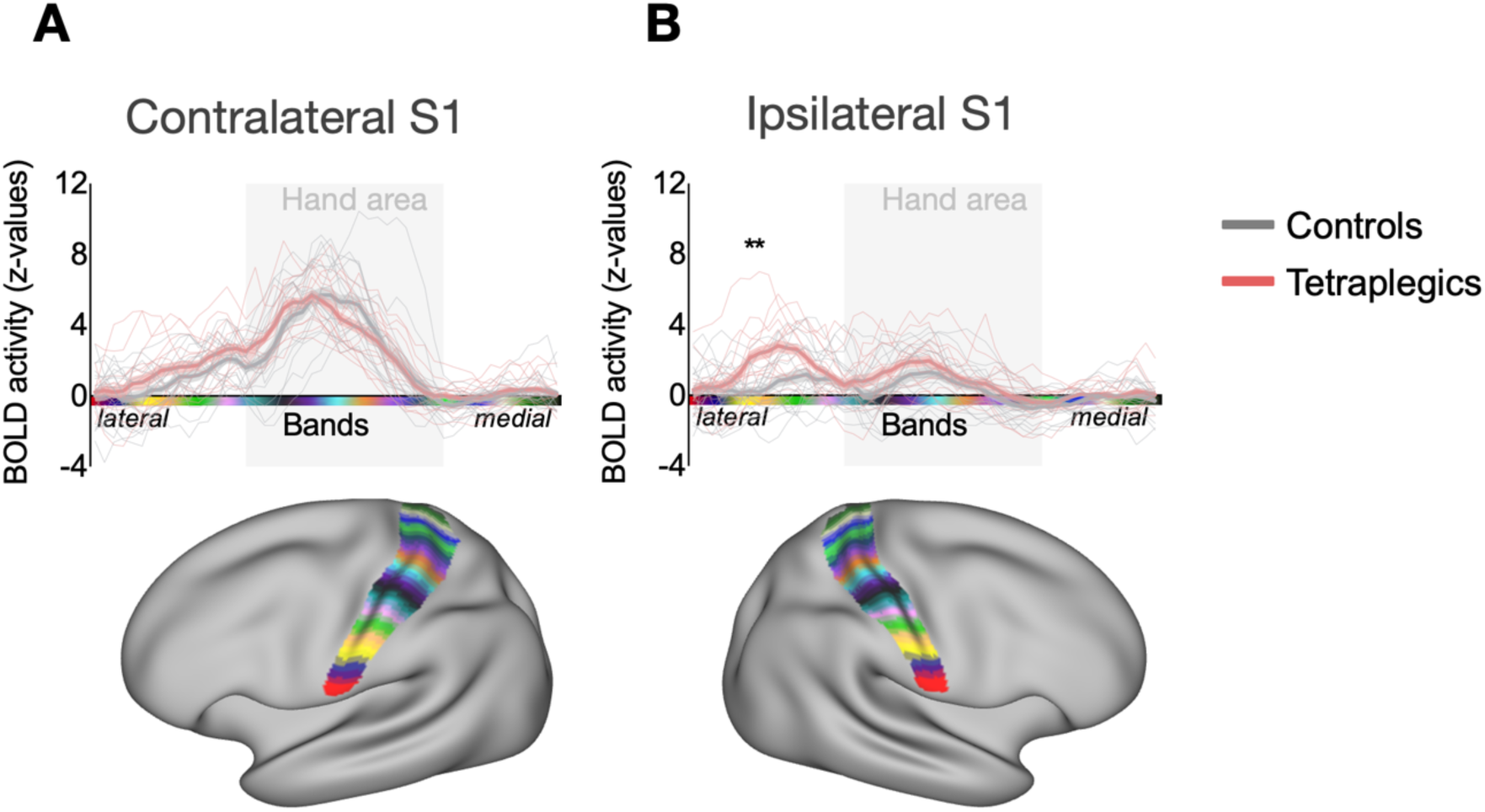
Band-wise fingers movement-related activation profiles in bilateral S1 during (attempted) finger tapping movements in individuals with tetraplegia and controls. A Finger movement-related activity (z-values) across 50 bands spanning contralateral S1 plotted for controls (gray) and individuals with tetraplegia (red). The shaded vertical area denotes the S1 bands within the S1 hand area (spanning 2cm above and below the anatomical hand knob). Both groups exhibit a pronounced peak in averaged finger movement-related activity within the contralateral S1 hand area. There was no significant difference in activation profile between groups. B Controls showed low finger-movement-related activation across ipsilateral S1, whereas individuals with tetraplegia displayed elevated activity with maximal group differences in bands lateral to the S1 hand area (significant cluster of elevated activity at band 7-8, as highlighted by asterisks). Error ribbons represent ±1 SEM across participants.

In ipsilateral S1, controls showed relatively low finger movement-related activity across ipsilateral S1 (**Fig. 1**), whereas individuals with tetraplegia exhibited qualitatively more elevated finger movement-related activity, most prominently in the S1 bands lateral to the hand representation. Comparison of the continuous spatial profiles identified a significant cluster of greater activity in individuals with tetraplegia lateral to the S1 hand area, spanning bands 7 to 8 (peak at band 8, Max z = 3.46; cluster p = 0.010, non-parametric permutation test corrected for multiple comparisons across all 50 bands).

When averaged across individuals with tetraplegia, finger movement-related activity was significantly greater than zero in both the ipsilateral and contralateral S1 hand areas. (**Appx. A2**; ipsilateral: t_(13)_ = 4.36, p < 0.01, BF_10_ = 49.05; contralateral: t_(13)_ = 9.16, p < 0.001, BF_10_ > 100). Controls showed robust activity in their contralateral S1 hand area, whereas evidence for recruitment of the ipsilateral S1 hand area was weak (ipsilateral: t_(17)_ = 1.96, p_corr_ = 0.07, BF_10_ = 1.16; contralateral: t_(17)_ = 9.89, p < 0.001, BF_10_ > 100;). Across hemispheres, there was no evidence for a general group difference in S1 hand-area activity (F_(1,21.66)_ = 1.07, p = 0.31, η^2^ = 0.02). As expected, activity was lower in ipsilateral than contralateral S1 in both groups (F_(1,15.38)_ = 201.06, p < 0.001, η^2^ = 0.45), and the degree of lateralization did not differ significantly between them (i.e., no interaction effect; F_(1,15.38)_ = 2.72, p = 0.11, η^2^ = 0.01).

Similar results were found for the contralateral primary motor cortex (M1) hand region of interest (ROI) (**Appx. A3**).

Together, these results indicate that while contralateral S1 activation remains preserved at normal levels, individuals with tetraplegia exhibit an increase in activity within ipsilateral S1 compared to controls. Notably, this increase in ipsilateral S1 activity was spatially localized lateral to the hand representation.

### IPSILATERAL S1 REPRESENTATIONAL STRUCTURE IS PRESERVED AFTER TETRAPLEGIA

We next used representational similarity analysis (RSA) to examine the multivariate organization of finger-specific activity patterns in the ipsilateral S1 hand area. We addressed three questions in sequence. First, whether the activity patterns associated with different fingers were reliably distinguishable on average. Second, whether the relative distances among individual finger pairs formed the expected representational geometry. Third, whether each participant’s entire pattern of pairwise distances resembled the canonical geometry of an able-bodied hand.

To address the first question, we examined how deafferentation affects the finger-related representational architecture in ipsilateral S1. We investigated the fine-grained inter-finger representational patterns in the ipsi- and contralateral S1 hand area of individuals with tetraplegia and controls using RSA.

Next, to quantify the separability of finger-specific activation patterns in ipsilateral S1, we calculated each participant’s mean cross-validated inter-finger distance, with larger values indicating more distinguishable finger-specific activity patterns. We found that the average inter-finger distance (**Fig. 2C**) was significantly greater than 0 in the ipsilateral S1 hand area for both controls (t_(17)_ = 13.96, p < 0.001) and individuals with tetraplegia (t_(12)_ = 8.64, p < 0.001). Furthermore, this average inter-finger distance in the ipsilateral S1 hand area was greater than in a cerebral spinal fluid (CSF) control ROI, where one would not expect finger-specific representational information, in both controls (t_(17)_ = 13.12, p < 0.01) and individuals with tetraplegia (t_(12)_ = 6.38, p < 0.01). Finally, the average inter-finger distance in the ipsilateral hemisphere was not significantly different between the control and tetraplegia groups (t(_29_) = -0.82, p = 0.42, BF_10_ = 0.44). Together, these findings indicate that finger movements evoke distinct activity patterns within the ipsilateral S1 hand area in both groups.

**Fig. 2.**
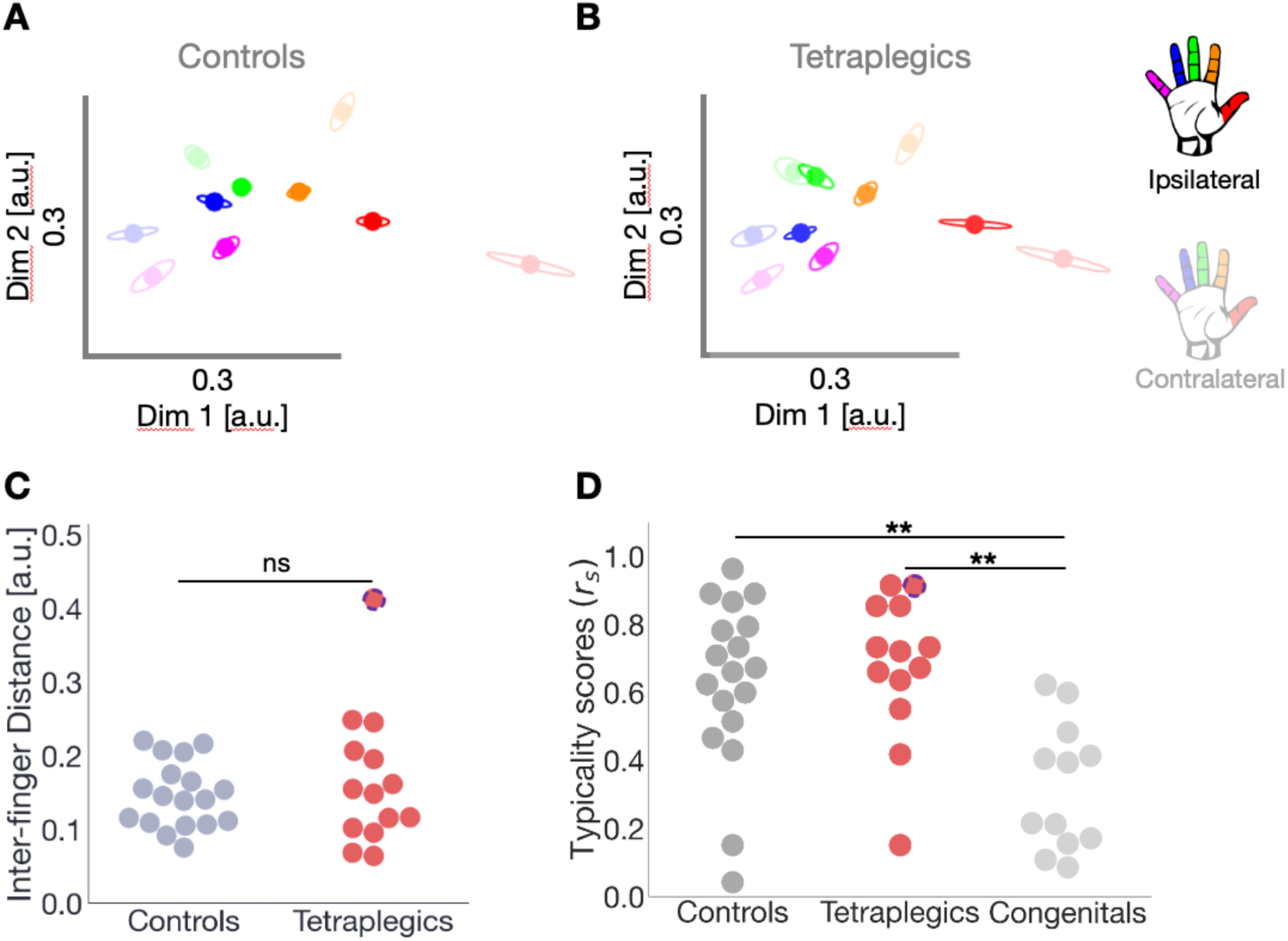
Finger somatotopy is preserved in the ipsilateral S1 hand area after tetraplegia. Two-dimensional visualization of the representational inter-finger distances in the ipsilateral S1 hand area for controls (**A**) and tetraplegic individuals (**B**). Individual fingers are represented by different colors: thumb (red); index finger (yellow); middle finger (green); ring finger (blue); little finger (pink). The relative distances between the dots reflect the inter-finger distances. Ellipses represent the between-participants’ standard error after Procrustes alignment. Opaque dots and ellipses reflect the fingers within the ipsilateral S1 hand area, while the more transparent dots reflect the fingers in the contralateral S1 hand area^33^. Dim = dimension; a.u. = arbitrary unit. **C** Mean cross-validated inter-finger distance was used to quantify overall finger-pattern separability in the ipsilateral S1 hand area for both controls (grey) and tetraplegic individuals (red). There was no significant difference in averaged inter- fingers distance between groups. The patient with no remaining sensorimotor function and any spared tissue bridges is highlighted with a blue circle. **D** Typicality of the representational structure in the ipsilateral S1 hand area in controls and individuals with tetraplegia, calculated as the spearman correlation coefficient (r_s_) between each participant’s ipsilateral representational geometry of fingers and a canonical RDM (defined as the control group’s average RDM from the contralateral S1 hand area). Congenital one-handers typicality scores (light grey) obtained from a previous study^37^ were added as an additional control to provide a biological baseline for the complete absence of a cortical hand representation. Controls and individuals with tetraplegia show higher typicality than congenital one-handers. There was no significant difference between controls and individuals with tetraplegia.

Qualitative inspection of the averaged inter-finger distance structure for both groups revealed characteristic inter-finger patterns in the ipsilateral and contralateral S1 hand areas (**Fig. 2A- B**)^1,33,36,38–40^. Neighboring fingers tended to have relatively smaller representational distances (i.e., more similar activation patterns) compared to those further apart. Furthermore, fingers used more frequently together in everyday life ^40^, such as the middle and ring fingers, appeared to have smaller representational distances compared to fingers that are used more independently, such as the thumb and index fingers.

Although the average inter-finger distance in the ipsilateral S1 hand area was comparable between individuals with tetraplegia and controls, differences in the relative distances between individual finger pairs could still be present, indicating specific alterations in representational geometry following tetraplegia. To address this, we first examined the inter-finger distances across pairs of fingers and between groups. Our mixed ANOVA revealed no significant difference in inter-finger distances between individuals with tetraplegia and controls (F_(1,30)_ = 0.67, p = 0.42, η^2^ = 0.01). The inter-finger distances were significantly different across finger pairs, as would be expected based on somatotopic mapping (F_(9,270)_ = 27.92, p < 0.001, η^2^ = 0.23). Importantly, this pattern of inter-finger distances in the ipsilateral S1 hand area was not significantly different between groups (i.e., no significant finger pair by group interaction; F_(9,270)_ = 0.91, p = 0.51, η^2^ = 0.01). Bayesian analysis of finger pair discriminability revealed anecdotal evidence for the null hypothesis (no difference between tetraplegic group and controls) for all ten finger pairs, with Bayes factors (BF_10_) ranging from 0.345 to 0.812.

While the pairwise analysis examined each inter-finger distance separately, we next asked whether each participant’s overall representational geometry resembled the canonical organization observed in controls. We calculated, per participant, a measure of representational typicality, or ‘normality’ in the ipsilateral S1 hand area. To do so, we performed a spearman correlation between each participant’s inter-finger distance pattern in the ipsilateral S1 hand area and a canonical inter-finger distance pattern (taken from the averaged group inter-finger distance pattern in the contralateral S1 hand area of the control group). We then compared the resulting ipsilateral S1 typicality scores between controls, tetraplegics, and an additional control group of congenital one-handers^36^ (**Fig. 2D**). The latter group consisted of individuals who were born without a hand and consequently do not have a representation of their missing hand. The typicality scores obtained from the S1 hand area contralateral to their missing hand were therefore included in the current study to have a proxy of ‘an absent hand representation’. Our ANOVA results suggested a significant difference in typicality between the three groups (F_(2,39)_ = 11.67, p < 0.001, η^2^ = 0.36). Ipsilateral S1 hand representation typicality of both individuals with tetraplegia and controls was significantly higher than the typicality scores of the congenital one-handers (controls: t_(17)_ = -4.12, p < 0.01, BF_10_ = 85.63; tetraplegics: t_(13)_ = -4.34, p < 0.01, BF10 = 105.05). Furthermore, ipsilateral S1 hand representation typicality of individuals with tetraplegia did not differ significantly from controls (t_(29)_ = -0.56, p = 0.83, BF_10_ = 0.39), with anecdotal evidence in favor of the null hypothesis.

### INTERHEMISPHERIC SIMILARITY OF S1 REPRESENTATIONAL STRUCTURES REMAIN HIGH AFTER TETRAPLEGIA

Because typical hand representations in healthy individuals are highly mirror-symmetric^6^, we next assessed whether the interhemispheric correspondence remains preserved after SCI, even if underlying interhemispheric excitability and inhibition may be altered. To do this, we quantified the interhemispheric similarity of the finger representational geometry between ipsilateral and contralateral S1 (**Appx. A4**). Using this approach, we found that interhemispheric similarity was generally high in both groups (Mantel r, controls: median 0.67; tetraplegics: median 0.72), and did not differ significantly between groups (t = 0.55, p = 0.59, BF10 = 0.39, i.e. anecdotal evidence in support of the null hypothesis).

### IPSILATERAL FINGER REPRESENTATIONS PERSIST WITHOUT ANY MEASURABLE PERIPHERAL INPUT

To directly test whether ongoing bottom-up sensory input is required to elicit ipsilateral finger representations, we next examined S01, the participant with complete paralysis of the tested hand, as previously confirmed using EMG ^41^, no measurable sensory function in that hand, and no detectable midsagittal tissue bridges. This participant also had no motor function and only minimal residual sensory function in the other hand (**Table 1**). Compared with the able-bodied control group, S01 exhibited a significantly greater mean inter-finger distance (FDR-corrected Crawford-Howell single-case comparison: t_(17)_ = 5.83, p < 0.001), indicating that the finger-specific activity patterns remained clearly distinguishable rather than being degraded. S01’s representational typicality was also significantly greater than that of congenital one-handers (t_(11)_ = 3.03, p < 0.05, BF_10_ = 5.3) and did not differ significantly from that of able-bodied controls (t_(16)_ = 1.15, p = 0.27, BF_10_ = 0.44). Although the latter comparison does not establish statistical equivalence, the combination of distinct finger-specific patterns and a typical representational geometry provide evidence that measurable bottom-up sensory input is not required to elicit an ipsilateral finger representation during attempted movement. Centrally generated movement-related signals are therefore sufficient to engage ipsilateral S1 somatotopically. Finally, exploring the individual with a clinically complete SCI separately revealed interhemispheric similarity did not differ significantly from controls (t_(17)_ = 1.12, p = 0.28) as evidenced again by a Crawford Howell test. Thus, interhemispheric correspondence of the S1 representational architecture is retained after tetraplegia, especially in that individual with no detectable connection between the hand and the brain.

### IPSILATERAL S1 MEASURES SHOW NO ROBUST ASSOCIATIONS WITH CLINICAL CHARACTERISTICS

To explore whether ipsilateral S1 measures were associated with clinical characteristics, we calculated bivariate correlations between average BOLD response or representational typicality and four clinical measures: years since injury, overall sensory function across both hands, overall motor function across both hands, and midsagittal tissue-bridge width. Motor function was included because the fMRI task involved executed or attempted finger movements and residual motor capacity could therefore contribute to the measured S1 responses. Pearson correlations were used for average BOLD response, whereas Spearman rank correlations were used for representational typicality. All reported p-values were adjusted for multiple comparisons using false-discovery-rate correction. Average BOLD response in the ipsilateral S1 hand area was not significantly associated with years since injury (r = -0.14, p_FDR_ = 0.63), overall sensory function (r = -0.55, p_FDR_ = 0.12), overall motor function (r = -0.12, p_FDR_ = 0.69), or midsagittal tissue-bridge width (r = -0.15, p_FDR_ = 0.63; **Fig. 3A-D**). The largest observed association was the negative correlation with sensory function, but this did not remain significant after FDR correction. Representational typicality was also not significantly associated with years since injury (r_s_ = -0.44, p_FDR_ = 0.36), overall sensory function (r_s_ = 0.17, p FDR = 0.79), overall motor function (r_s_ = 0.28, p_FDR_ = 0.67), or midsagittal tissue-bridge width (r_s_ = -0.09, p_FDR_ = 0.79; **Fig. 3E-H**). Given the small sample and exploratory nature of these analyses, the non-significant findings should not be interpreted as evidence that ipsilateral S1 activity or representational typicality is unrelated to these clinical characteristics.

**Fig. 3.**
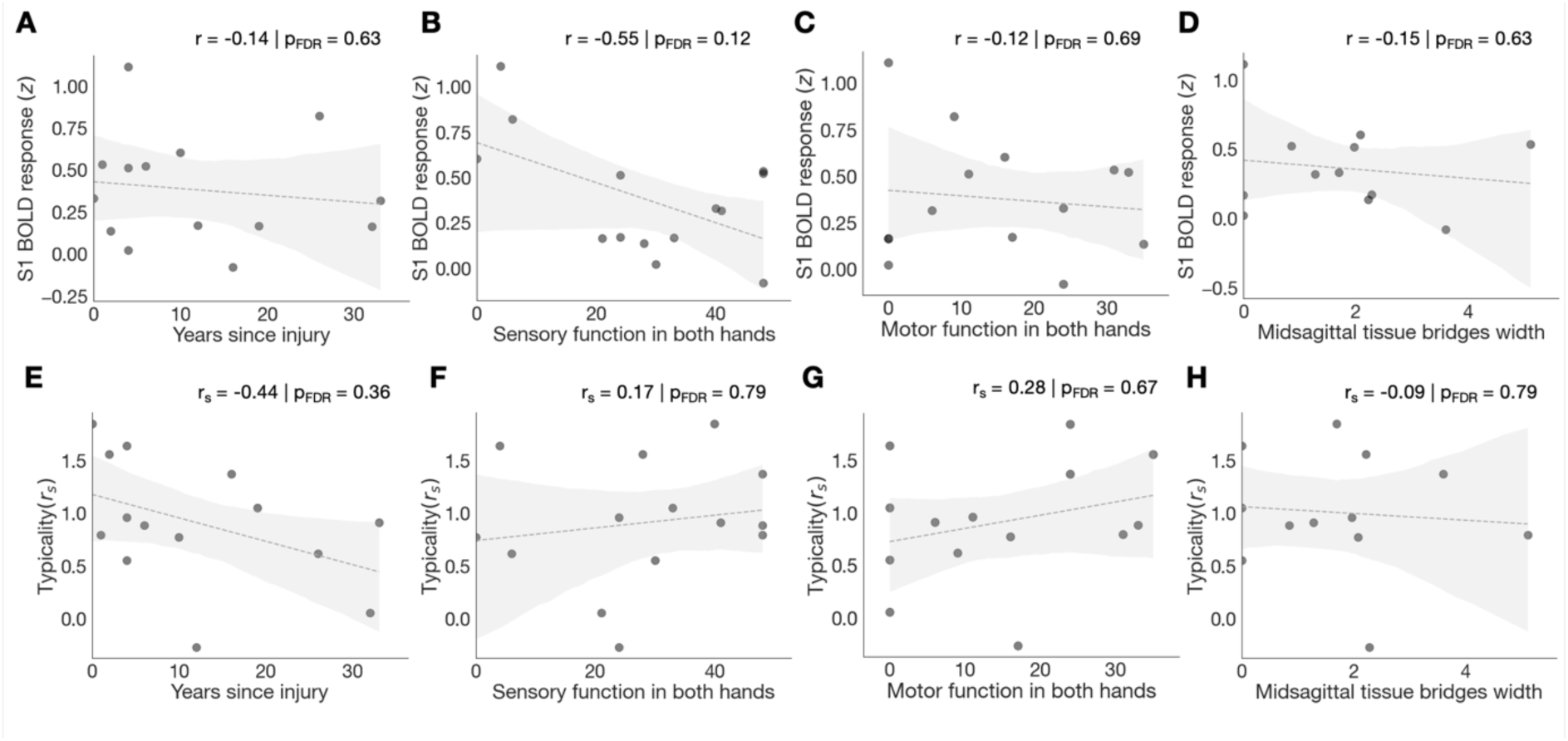
Associations between clinical characteristics and ipsilateral S1 measures. Scatterplots show the relationships between four clinical characteristics and average BOLD response magnitude (top row; **A-D**) or representational typicality (bottom row; **E-H**) within the ipsilateral S1 hand area of individuals with tetraplegia. The clinical measures are years since injury (**A, E**), overall sensory function across both hands (B, F), overall motor function across both hands (**C, G**), and midsagittal tissue- bridge width (**D, H**). Pearson correlation coefficients (r) are reported for average BOLD response, and Spearman rank-correlation coefficients (r_s_) for representational typicality. Values above each panel show the correlation coefficient and the false-discovery-rate-adjusted p-value (p_FDR_). Dashed lines indicate the linear line of best fit, shaded areas denote the corresponding 95% confidence intervals, and dots represent individual participants. Sample size (n = 12) was lower for D and H due to data availability. None of the associations remained significant after FDR correction.

To more comprehensively evaluate the combined predictive value of these clinical metrics on ipsilateral S1 activity, we computed robust multiple linear regression models predicting either average ipsilateral S1 hand area activity or representational typicality from sensory function, motor function, tissue bridge width, and years since injury. In the model predicting mean ipsilateral S1 BOLD response, none of the clinical predictors showed a statistically significant independent association with activation magnitude. In the model of average ipsilateral S1 BOLD response, none of the four clinical variables showed a statistically significant independent association with activity magnitude. Sensory function showed a negative but non-significant coefficient (b = -0.0097, SE = 0.0077, z = -1.26, p = 0.206, 95% CI [- 0.025,0.005] [-0.025,0.005]). Motor function was not associated with activity (b=0.00008, SE = 0.012, z = 0.01, p = 0.995, 95% CI [-0.024,0.024] [-0.024,0.024]), nor were midsagittal tissue-bridge width (b = -0.0030, SE = 0.090, z = -0.03, p = 0.974, 95% CI [-0.179,0.174] [- 0.179,0.174]) or years since injury (b = -0.0067, SE = 0.013, z = -0.54, p = 0.592, 95% CI [- 0.031,0.018] [−0.031,0.018]). Similarly, none of the clinical variables showed a statistically significant independent association with representational typicality. The coefficients were *b* = 0.0011 b = 0.0011 for sensory function (SE = 0.015, z = 0.08, p = 0.940, 95% CI [- 0.027,0.030 ] [-0.027,0.030]), b = 0.0101 for motor function (SE = 0.023, z = 0.43, p = 0.668, 95% CI [- 0.036,0.056 ] [-0.036,0.056]), b = -0.0930 for midsagittal tissue-bridge width (SE = 0.171, z = -0.55, p = 0.586, 95% CI [-0.427,0.241] [-0.427,0.241]), and b = -0.0092 for years since injury (SE = 0.024, z = -0.39, p = 0.698, 95% CI [-0.056,0.037 ] [-0.056,0.037]).

Thus, none of the four clinical variables showed a statistically significant association with either ipsilateral S1 outcome in the bivariate analyses or a statistically significant independent coefficient in the robust regression models.

## DISCUSSION

We investigated whether finger-specific representations in ipsilateral S1 can be engaged without the peripheral signals that normally accompany hand movement. We found that ipsilateral S1 hand representations are preserved and can be activated by attempted finger movements in individuals with tetraplegia. Crucially, finger representational typicality in the ipsilateral S1 hand area did not differ significantly from that of healthy controls, and a typical representation was evident in a participant with a complete loss of sensorimotor hand function and a complete disconnect between the brain and periphery (i.e. no detectable spared spinal tissue bridges). Because this preserved topographic organization persisted in the absence of peripheral hand-related inputs, these findings provide direct evidence that ipsilateral S1 representations can be driven entirely by central, top-down mechanisms.

This interpretation is consistent with previous evidence that ipsilateral representations are strongly shaped by task context and active movement ^1,4–8,15,18,19,35,42^, and extends it by showing that hand-related peripheral input is not required for their expression. Indeed, previous work implicated centrally generated processes in ipsilateral S1 activity^1,19,43^, but could not establish whether these signals were sufficient because hand-related peripheral input was never fully excluded. Our findings close this gap by showing that a typical finger-specific ipsilateral representation can be elicited during attempted movement despite complete loss of sensorimotor hand function and no detectable midsagittal tissue bridges. This advances the interpretation of ipsilateral representations from central contribution to central sufficiency. We note that this inference rests on one fully characterized individual, supported by group-level equivalence of typicality, and would benefit from replication in further individuals with clinically complete injuries.

The precise nature of central signals engaging ipsilateral S1 representations remains uncertain and involves several interacting sources. Given that our participants performed attempted movements, efference copies or corollary-discharge signals conveying the predicted sensory consequences of movement are a prime candidate. However, S1 is not merely a passive recipient of motor or sensory signals. It is sensitive to broader cognitive modulation. Previous studies have demonstrated that contralateral S1 can be activated in a somatotopically specific manner through cognitive processes, including finger-specific attention^44^, touch expectation^45^, motor imagery^46,47^, working memory^48^, and action observation^49^. Our findings extend this framework to ipsilateral S1, showing that centrally generated influences are sufficient to active its fine-grained finger representation even after severe and longstanding disruption of sensorimotor communication with the hand. These signals may reach ipsilateral S1 through transcallosal interactions with the opposite sensorimotor cortex or through inputs from higher-order motor areas with bilateral organization.

To characterize the spatial distribution of movement-related activity across S1, we divided the cortical S1 ribbon into 50 equidistant bands running from lateral to medial and compared the activation profiles between groups. This analysis revealed broadly elevated ipsilateral BOLD responses in tetraplegic individuals, which peaked laterally to the canonical hand representation. Although group differences within the anatomically defined hand area itself did not reach statistical significance, potentially due to limited statistical power, the overall pattern of elevated ipsilateral recruitment suggests that the loss of ascending peripheral input may trigger broad network-level adjustments. A coexistence of altered movement-related activity and preserved representational organization is consistent with a homeostatic account of deprivation-related plasticity^50^. Following the loss of its usual sensory drive, the cortex may compensate by adjusting local excitability and inhibition, thereby increasing its responsiveness to inputs that were already present. Under this account, deprivation may change how strongly finger-related signals are expressed, without necessarily changing the information encoded by the underlying population. A related dissociation has been reported after acquired hand loss, increasing motor demands during intact-hand movements elicited greater activity in the deprived ipsilateral S1 hand area without a corresponding change in finger representational organization^51^ . At the circuit level, altered interhemispheric inhibition could provide one route through which pre-existing bilateral movement-related signals become more strongly expressed after deafferentation^52^. During unilateral movement, contralateral S1 can receive movement-related signals even when proprioceptive feedback is absent, potentially through interactions with premotor cortex^53,54^. These findings establish that S1 receives centrally generated movement-related input, but they neither identify this input specifically as an efference copy nor determine whether the same mechanism drives ipsilateral S1 activity. Ipsilateral S1 could receive movement-related information through transcallosal interactions with the contralateral sensorimotor cortex or through projections from higher-order motor areas with bilateral organisation^6,15,42,43^. One possibility is that deafferentation increases the gain of these pre-existing central inputs. Acute deafferentation has been shown to alter cortical excitability and interhemispheric interactions^52^. However, this evidence primarily concerns M1 and cannot be assumed to generalize directly to S1. More generally, homeostatic mechanisms could increase cortical responsiveness following the loss of peripheral input without requiring a change in the underlying somatotopic architecture^50,55^. We therefore regard homeostatic amplification of central inputs including, potentially, transcallosal inputs as a plausible but untested explanation for the elevated ipsilateral S1 activity observed after tetraplegia. The present fMRI data cannot determine whether this effect reflects altered excitation, inhibition, or both.

We did not detect significant associations between ipsilateral S1 activity or representational typicality and the examined clinical characteristics. These exploratory analyses should however be interpreted with caution. While our sample size (n=14) is typical for neuroimaging studies in specialized SCI populations ^26,56,57^, its relatively small size limited our power to detect subtle effects, particularly in the correlational analyses involving multiple predictors. The absence of significant associations thereby does not establish that ipsilateral S1 activity or representational organization is independent of injury severity, residual spinal tissue, or time since injury. Importantly, however, finger-specific ipsilateral representations were detectable across individuals with widely varying clinical profiles, including those with severe sensorimotor impairment. Their expression is therefore not confined to individuals with substantial residual hand function.

Together, our findings show that centrally generated movement-related signals are sufficient to engage finger-specific representations in ipsilateral S1 after tetraplegia, even without overt movement or measurable hand-related sensory input. More broadly, they indicate that profound sensorimotor disconnection does not render ipsilateral S1 functionally inaccessible but leaves an established finger-specific representation available to central motor processes. This persistence may provide additional cortical signals or targets for movement decoding, artificial sensory feedback, and closed-loop neuroprosthetic integration. Ipsilateral S1 may be particularly relevant when conventional contralateral targets are compromised or surgically inaccessible, offering a complementary substrate through which residual sensorimotor organization could be accessed. However, the causal and practical contribution of ipsilateral S1 to percept generation and prosthetic control remains to be established.

## METHODS

The data used in this manuscript have been previously published in eLife^36,58^. In the previous manuscript we used this dataset to examine preserved contralateral representations in S1. Here we focus on their ipsilateral counterpart. The experimental task, fMRI data acquisition, fMRI data preprocessing, and behavioral and clinical assessments are therefore identical to what was described in^58^. We briefly restate them here for the reader’s convenience.

### PARTICIPANTS

We recruited 15 chronic tetraplegic participants, of which 14 completed the measurements (mean age ± s.e.m. = 55 ± 3.6 years; one female; six dominant left-handers; see **Table 1** for demographic and clinical details). The following inclusion criteria applied to our recruitment: aged 18–75 years, no MRI contraindications, chronic tetraplegia (i.e. > 6 months post injury), no neurological impairment or body function impairments not induced by SCI, and able to provide informed consent.

We further recruited 18 age-, sex-, and handedness-matched able-bodied control participants (age = 56 ± 3.6 years; one female; five dominant left-handers). Inclusion criteria for the control participants included: aged 18-75 years, no MRI contraindications, no impairment of body function induced by SCI, no neurological illness, no hand impairments, and able to provide informed consent.

We obtained participants’ informed consent according to the Declaration of Helsinki prior to study onset. The same was previously done for the congenitals^36^. Ethical approval was granted by the Kantonale Ethikkommission Zürich (KEK-2018-00937) and this study is registered on clinicaltrials.gov under NCT03772548. Two individuals with tetraplegia and one control participant were rescanned due to excessive head motion during fMRI acquisition or suboptimal slice placement. We had to exclude one tetraplegic individual due to withdrawal from the study and one control participant due to distorted data.

### CLINICAL CHARACTERISATION

We collected clinical data in a separate session, assessing individuals with tetraplegia’s completeness of injury and impairment level using the International Standards for Neurological Classification of Spinal Cord Injury (ISNCSCI). To evaluate residual sensorimotor upper limb function, we gathered GRASSP scores ^24^, where the maximum score per upper limb is 116, indicating healthy function. For the robust linear regression models investigating ipsilateral S1 activity and typicality, motor GRASSP scores were added across ’more-impaired hand’ and the ’less-impaired hand’ for each tetraplegic individual. The ’more-impaired hand’ was defined as the hand participants were instructed to move during the fMRI task (referred to as ’Hand tested’ in **Table 1**), and its motor GRASSP score was the sum of its relevant motor sub-components as listed in **Table 1** (column ’GRASSP tested Hand. motor/sensory’, motor component). Similarly, the ’less-impaired hand’ was the stationary hand (referred to as ’GRASSP other Hand.’ in **Table 1**), and its motor GRASSP score was the sum of its motor sub-components as listed in Table 1 (column ’GRASSP other Hand. motor/sensory’, motor component).

### EXPERIMENTAL PROCEDURE AND TASKS

To explore fine-grained somatotopic representations ipsilateral to the moved fingers, we used 3 tesla functional MRI. Participants were instructed to make unimanual individual fingers movements while their palm was facing upwards. We only tested individuals with tetraplegia’s most impaired upper limp (determined using the GRASSP score). For some tetraplegic individuals the loss of sensorimotor function after SCI did not allow them to make overt movements. They were then carefully instructed to make attempted (i.e., not imagined) movements. Our fMRI paradigm was carried out in a blocked design fashion with six conditions: movement conditions for each of the five fingers and a rest condition. Participants viewed a screen with five horizontally aligned white circles corresponding to the five fingers. For participants moving their left hand the leftmost and rightmost circles corresponded to the thumb and little finger, respectively. For participants moving the right hand the leftmost circle corresponded to the little finger and the rightmost circle to the thumb. To instruct participants to make self-paced flexion/extension with one of the fingers, the corresponding circle on the screen turned red. The word ‘Rest’ on the screen indicated a rest condition during which participants were instructed to not move. A movement or rest block lasted 8s, and each condition was repeated five times per run in a counterbalanced order. Each run comprised a different block order and had a duration of 4 min and 14 s. We acquired four runs, with a total duration of 16 min and 56 s. Instructions were delivered using Psychtoolbox (v3) implemented in MATLAB (v2014). We minimized head motion using over-ear MRI-safe headphones or padded cushions.

### MRI DATA ACQUISITION

We acquired MRI data using a Philips 3 Tesla Ingenia system (Best, The Netherlands) with a 17-channel HeadNeckSpine or, in case of participant discomfort due to the coil’s narrowness, a 15-channel HeadSpine coil. Anatomical T1-weighted images covering the brain and cervical spinal cord were acquired using the following acquisition parameters: 0.8mm^3^ resolution, repetition time (TR) = 9.3ms, echo time (TE) = 4.4ms, and flip angle 8°. Sagittal T2-weighted anatomical images of the cervical spinal cord were acquired with a resolution of 1 × 1 × 3 mm. The acquisition used a repetition time (TR) of 4500 ms, an echo time (TE) of 85 ms, a flip angle of 90°, and a slice gap of 0.3 mm, covering 15 slices. Task-fMRI data were acquired using an echo-planar-imaging (EPI) sequence with partial brain coverage: 22 sagittal slices were centered on the anatomical location of the hand knob with coverage over the thalamus and brainstem. We used the following acquisition parameters: 2 mm^3^ resolution, TR = 2000ms, TE = 30ms, flip angle = 82°, and SENSE factor = 2.2. We acquired 127 volumes for each blocked design run.

### ANALYSIS OF FMRI DATA

fMRI analysis was implemented using FSL v6.0 (https://fsl.fmrib.ox.ac.uk/fsl/fslwiki), Advanced Normalization Tools (ANTs) v2.3.1 (http://stnava.github.io/ANTs), the RSA toolbox ^59,60^, and MATLAB (R2018a).

### QUANTIFICATION OF MIDSAGITTAL TISSUE BRIDGES

We quantified midsagittal tissue bridges on sagittal T2-weighted images of the cervical spinal cord at the lesion level using Jim 7.0 software. Following established procedures ^61–63^, an experimenter blinded to tetraplegics’ identity performed manual lesion segmentation on the midsagittal slice, including only scans where the lesion was clearly visible. Tissue bridges were defined as the relatively hypointense intramedullary region spanning the lesion, and the total width was calculated as the sum of ventral and dorsal tissue-bridge widths. Images from two individuals with tetraplegia were excluded due to metal artifacts or insufficient data quality, which prevented reliable lesion measurement.

### PREPROCESSING OF FMRI DATA

Common preprocessing steps were applied using FSL’s Expert Analysis Tool (FEAT). The following preprocessing steps were included: motion correction using MCFLIRT ^64^, brain extraction using automated brain extraction tool BET ^65^, spatial smoothing using a 2mm full-width-at-half-maximum (FWHM) Gaussian kernel, and high-pass temporal filtering with a 100s cut-off.

Image co-registration was done in separate, visually inspected, steps. For each participant, a midspace was calculated between the four blocked design runs, that is, an average space in which images are minimally reoriented. We then transformed all fMRI data to this midspace using purely rigid probability mapping in ANTs. Next, we registered each participant’s midspace to the T1-weighted image, initially using 6 degrees of freedom and the mutual information cost function, and then optimized using boundary-based registration (BBR; ^66^) Each co-registration step was visually inspected and, if needed, manually optimized using blink comparison in Freeview.

### REGIONS OF INTEREST DEFINITION

For the univariate and multivariate analyses focused on the hand representation, a participant-specific S1 hand ROI was generated in each participant’s native anatomical space using FreeSurfer (recon-all) and Brodmann-area labels. Specifically, the ROI was defined as the union of BA1, BA2, BA3a, and BA3b within S1, converted to a volumetric mask, with any holes filled and the resulting non-zero voxels mean-dilated to obtain a contiguous, analysis-ready ROI. To restrict this ROI to the hand region, we selected the axial slices spanning ±2 cm medial/lateral around the hand knob and retained only the corresponding S1 voxels within this slab ^67^. The same bilateral hand-region masks were applied to contralateral and ipsilateral hemispheres.

For the cortical band analysis, we instead used an anatomical S1 mask defined in MNI space (obtained via https://github.com/DiedrichsenLab/fs_LR_32 based on the Glasser et al. S1 parcellation^68^) in order to apply a consistent banding procedure across participants within a common coordinate system.

### UNIVARIATE ANALYSIS

To assess univariate task-related activity, time-series statistical analysis was carried out per run using FMRIB’s Improved Linear Model (FILM) with local autocorrelation correction, as implemented in FEAT. We obtained activity estimates using a general linear modelling (GLM) based on the double-gamma HRF and its temporal derivative. Each finger movement condition was contrasted with rest. A further contrast was defined for overall task-related activity by contrasting all movement conditions with rest. A fixed effects higher-level analysis was run for each participant to average across runs.

The z-scored BOLD response for overall task-related activity was then extracted for voxels underlying the contra- and ipsilateral S1 hand ROIs per participant. A similar analysis was used to investigate overall task-related activity in contra- and ipsilateral M1 hand ROIs (see **Appx A3**).

### S1 BANDS ANALYSIS

To quantify the distribution of average movement-related activity across contra- and ipsilateral S1 during movements, we segmented S1 into 50 equidistant bands perpendicular to the central sulcus (**Fig. 1**), following the approach described by Schone et al ^69^. The primary S1 hand area was anatomically defined by selecting the axial slices spanning ±2 cm medial/lateral around the hand knob within S1. Second-level statistical maps from the overall task-related activity, reflecting average movement-related activity, were projected onto each participant’s cortical surface using cortical-ribbon mapping and then aligned to the fs-LR surface template. We extracted average activation (z-value) in each of the 50 contra- and ipsilateral S1 bands. To analyze differences between individuals with tetraplegia and controls, we employed non-parametric permutation t-tests using the SPM1D Matlab toolbox, which treats band data as a continuous field and accounts for spatial correlations using random field theory. Group-level comparisons were conducted separately for the contralateral and ipsilateral hemispheres.

### REPRESENTATIONAL SIMILARITY ANALYSIS

The somatotopic hand representation is characterized by representational distances between fingers that has been shown to be consistent across control individuals in both the contra- ^40,70–75^ and ipsilateral sensorimotor cortex ^1,6^. Such intricate inter-finger relationship patterns can be estimated using representational similarity analysis (RSA^76^). In this study we therefore used RSA to estimate the intricate representational relationship between finger representations in the ipsilateral S1 hand area of both individuals with tetraplegia and controls. We computed the distance between the activity patterns measured for each finger pair within the ipsilateral S1 hand ROI using the cross-validated squared Mahalanobis distance (or crossnobis distance) ^77^. We extracted the blocked design voxel-wise parameter estimates (betas) for each finger movement condition versus rest and the model fit residuals under the S1 hand ROI. We prewhitened the extracted betas using the model fit residuals. We then calculated the cross-validated squared Mahalanobis distances between each possible finger pair, using our four runs as independent cross-validation folds, and averaged the resulting distances across the folds. If it is impossible to statistically differentiate between conditions (i.e., when this parameter is not represented in the ROI), then the expected value of the distance estimate would be 0. If it is possible to distinguish between activity patterns, then this value will be larger than 0. We assembled all finger pair distances in a representational dissimilarity matrix (RDM), with a width and height corresponding to the five finger movement conditions. Since the RDM is mirrored across the diagonal with meaningless zeros on the diagonal, all statistical analyses were conducted on the 10 unique off-diagonal values of the RDM. We first estimated the strength of the finger representation or ‘averaged inter-fingers distance’ by averaging the 10 unique off-diagonal values of the RDM. If there is no information in the ROI that can statistically distinguish between the finger conditions, then due to cross-validation the expected averaged inter-fingers distance would be 0. If there is differentiation between the finger conditions, the averaged inter-fingers distance would be larger than 0^77^. To further ensure that our S1 hand ROIs was activated distinctly for different fingers, we created a CSF ROI that would not contain finger-specific information. We repeated our RSA analysis in this ROI and statistically compared the averaged inter-fingers distance of the CSF and ipsilateral S1 hand area ROIs. Second, we tested whether the inter-finger distances were different between controls and individuals with tetraplegia using a robust mixed ANOVA with a within-participants factor for finger pair (10 levels) and a between-participants factor for group (two levels: controls and individuals with tetraplegia). Third, we estimated the somatotopic typicality (or normality) of each participant’s RDM for both individuals with tetraplegia and controls ^33,36,40,72,78^ by correlating it with a canonical RDM using a mantel permutation test. The canonical RDM was computed as the group average of the controls. To validate our typicality scores we also computed typicality scores using a canonical RDM from another study (only contralateral typicality data; publicly available on https://osf.io/gmvua/) ^36^. The resulting spearman correlations were Fisher r-to-z transformed prior to statistical analysis (the spearman correlations (r*_s_*) are used solely for visualization). Controls’ and individuals with tetraplegia’s correlations coefficients were compared to each other and to those of a group of individuals with congenital hand loss (n = 13), hereafter one-handers, obtained in another study (data publicly available on https://osf.io/gmvua/) ^36^. Congenital one-handers are born without a hand and do not have an S1 hand representation contralateral to the missing hand. Finally, we performed multidimensional scaling (MDS) to visualize the distance structure of the RDM in both contra- and ipsilateral S1 in an intuitive manner. MDS projects the higher-dimensional RDM into a lower-dimensional space, while preserving the inter-finger distance values as well as possible ^79^. MDS was performed for each individual participant and then averaged per group after Procrustes alignment to remove arbitrary rotation induced by MDS.

To quantify interhemispheric correspondence, we calculated a Mantel correlation between the 10 unique off-diagonal entries of each participant’s ipsilateral and contralateral S1 hand-area RDMs. The resulting Spearman correlation coefficients were Fisher r-to-z transformed before group-level analysis and compared between individuals with tetraplegia and controls using a two-sample test.

### CLINICAL ASSOCIATION ANALYSIS

Clinical association analyses We examined associations between ipsilateral S1 measures and four prespecified clinical variables: overall sensory function across both hands, overall motor function across both hands, midsagittal tissue-bridge width, and years since injury. Overall sensory and motor function were calculated by summing the corresponding GRASSP component scores across the tested and non-tested hands. The functional outcomes were the average movement-related BOLD response within the S1 hand area ipsilateral to the tested hand and Fisher r-to-z-transformed representational typicality within the same region. First, we performed exploratory bivariate analyses using all observations available for each association. Pearson correlations were used for average BOLD response, whereas Spearman rank correlations were used for representational typicality. The associated *p* p-values were adjusted for multiple comparisons using the Benjamini–Hochberg false-discovery-rate procedure across the eight bivariate correlations. Second, we fitted separate robust multiple linear regression models for average ipsilateral S1 activity and representational typicality. Each model included overall sensory function, overall motor function, midsagittal tissue- bridge width, and years since injury as simultaneous predictors. Because tissue-bridge measurements were unavailable for two participants, these models used complete cases and included 12 individuals. Robust linear models were fitted in statsmodels using iteratively reweighted least squares with Huber’s T norm. Model scale was estimated using the median absolute deviation, and H1 heteroscedasticity-robust covariance estimates were used to calculate standard errors, two-sided Wald z-tests, and 95% confidence intervals. The regression coefficients are unstandardized and therefore express the expected change in the respective ipsilateral S1 outcome for a one-unit increase in the predictor while holding the other three clinical variables constant. Given the limited sample size and four simultaneous predictors, the regression models were considered exploratory.

### STATISTICAL DATA ANALYSIS

Statistical analysis was carried out using statsmodels (v0.13.1). Standard approaches were used for statistical analysis, as mentioned in the Results section. To detect outliers, we used the robustbase toolbox ^80^. **S_n_** identifies an outlier (x_i)_ if the median distance of x_i_ from all other points, was greater than the outlier criterion (λ=2) times the median absolute distance of every point from every other point:

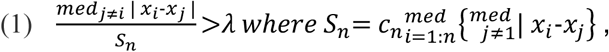

where c_n_ is a bias correction factor for finite sample sizes ^81^. We detected no outliers for clinical data and z-scored BOLD responses and one outlier for the averaged inter-fingers distance and typicality scores. These outliers were excluded from any further analysis.

For the band-wise analysis of S1 activation profiles, any identified outlier data points within a participant’s 49-band sequence were replaced with the mean value of their immediately adjacent non-outlier bands. This step was taken to ensure continuous and visually uninterrupted band plots for clearer interpretation, while minimally affecting the overall profile shape.

If normality was violated (assessed using the Shapiro–Wilk test), non-parametric statistical testing or robust ANOVAs were used trough the bioinfo-kit and pingouin toolbox ^82,83^. The assumption of sphericity was testing using Mauchly’s test of sphericity and if sphericity was violated, a Greenhouse-Geisser correction was applied. All testing was two-tailed, and corrected p-values were calculated using the Benjamini–Hochberg procedure to control the FDR with q < 0.05. For the band analysis, we conducted non-parametric two-sample t-tests at each of the 50 equidistant bands spanning S1. The resulting SPM{t} trajectory was thresholded based on random field theory (RFT) to control for multiple comparisons across the continuum of bands^84^. A critical statistic threshold (z*) was derived to maintain a family-wise error rate of p < 0.05, and the p-value for any supra-threshold clusters was reported as pset. The correlational analysis was considered exploratory.

Bayesian analysis was carried out using pingouin toolbox for our main comparisons to investigate support for the null hypothesis, i.e., no differences between groups, with a Cauchy prior width set at 0.707. Following the conventional cut-offs, a BF smaller than 1/3 is considered substantial evidence in favor of the null hypothesis. A BF greater than 3 is considered substantial evidence, and a BF greater than 10 is considered strong evidence in favor of the alternative hypothesis. A BF between 1/3 and 3 is considered weak or anecdotal evidence ^85,86^.

For single-case analyses, we used Crawford–Howell modified *t* t-tests to compare S01’s inter-finger distance, representational typicality, and Fisher-transformed interhemispheric similarity coefficient with the corresponding control distributions. This test treats the control sample as a sample rather than as a population and is therefore appropriate for comparing one individual with a small control group. The resulting p-values were corrected across the three prespecified S01 comparisons using the Benjamini–Hochberg false-discovery-rate procedure.

## ACHKNOWLEDGMENTS

We thank our participants for taking part in the study. We thank Michaela Verling, Lydia Kämpf, Nicolin Gauler, and Silvia Hofer for help with data collection. We thank Roger Lüchinger for technical support.

## SUPPLEMENTARY MATERIAL

## APPENDIX A1

To visualize the spatial extent of task-evoked activity, we computed group-level activation maps for the unimanual finger tapping contrast (Movement > Rest) for both control participants and individuals with tetraplegia. As shown in Figure A1, both groups engaged a widespread sensorimotor network. Significant activation clusters (Z > 3.1, cluster-corrected p < 0.05) were observed primarily in the contralateral primary sensorimotor cortices (S1/M1) but also extended bilaterally into ipsilateral sensorimotor regions, supplementary motor area (SMA), and secondary somatosensory cortex.

**Appx. A1.**
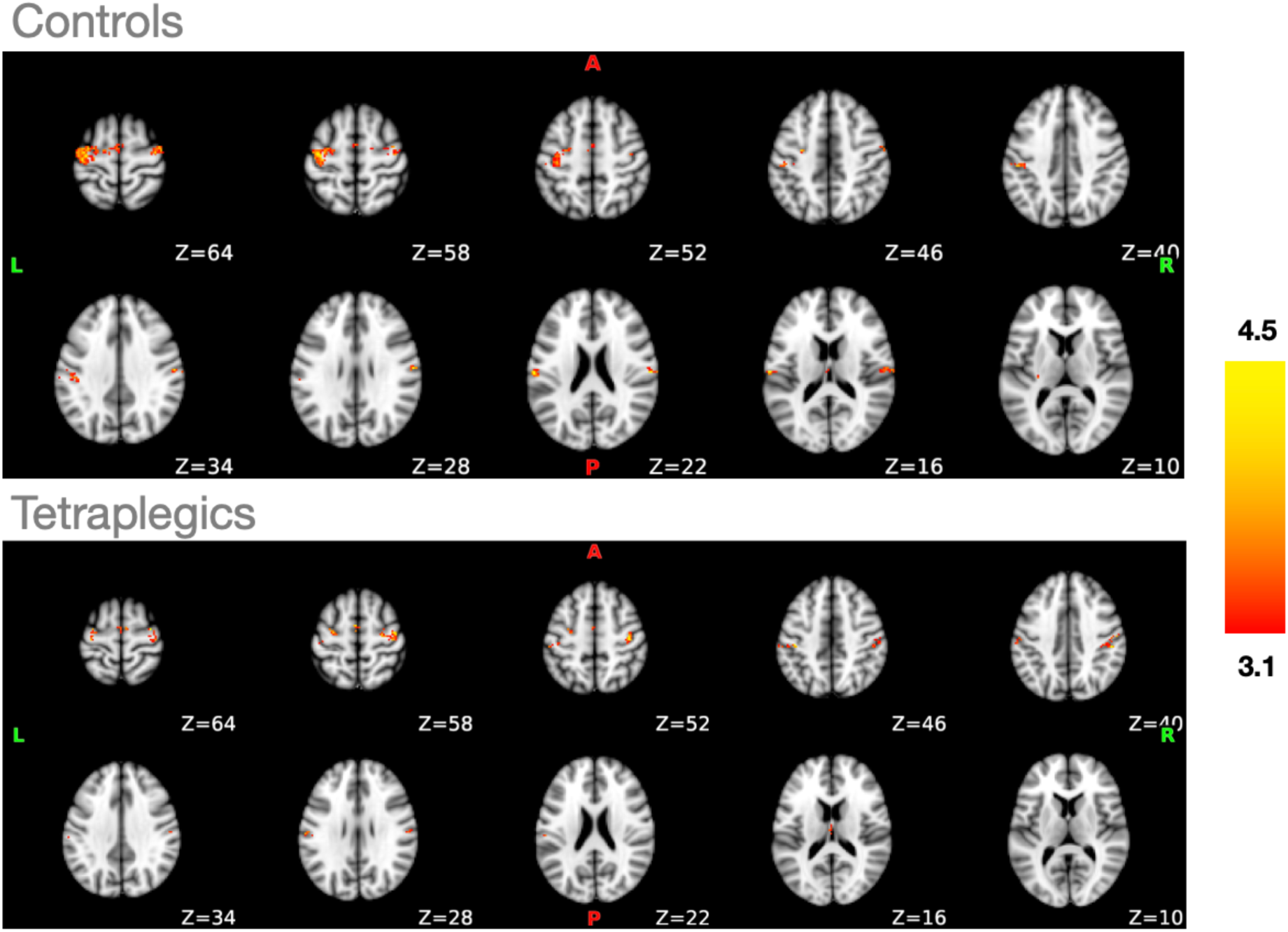
Group-level activation maps during unimanual finger movements. Axial slices showing significant BOLD activation (Z > 3.1, cluster-corrected p < 0.05) for the contrast Unimanual Finger Movement > Rest in control participants (top) and individuals with tetraplegia (bottom). Both groups show robust activation in the contralateral (left) primary sensorimotor cortex, ipsilateral (right) sensorimotor regions, bilateral supplementary motor cortex, and bilateral secondary somatosensory cortex. The color bar represents Z-statistics. L = Left hemisphere (contralateral to the moved hand); R = Right hemisphere (ipsilateral to the moved hand). Participants moved either hand, so some maps have been flipped to compare them on a group level.

## APPENDIX A2

Individuals with tetraplegia significantly activated both their ipsi- and contralateral S1 hand area through unimanual individual finger movements (ipsilateral: t_(13)_ = 4.36, p < 0.01, BF_10_ = 49.05; contralateral: t_(13)_ = 9.16, p < 0.001, BF_10_ > 100), while controls significantly activated their contralateral S1 hand area (t_(17)_ = 9.89, p < 0.001, BF_10_ > 100) and their ipsilateral S1 engagement trended towards significance (t_(17)_ = 1.96, p_corr_ = 0.07, BF_10_ = 1.16). A robust mixed ANOVA revealed no significant difference in task-related activity between controls and individuals with tetraplegia (F_(1,21.66)_ = 1.07, p = 0.31, η^2^ = 0.02). As expected, average movement-related activity levels were lower in the ipsi- than in the contralateral S1 hand area (F_(1,15.38)_ = 201.06, p < 0.001, η^2^ = 0.45). This difference between averaged contra- and ipsilateral movement-related activity levels was not significantly different between groups (i.e., no interaction effect; F_(1,15.38)_ = 2.72, p = 0.11, η^2^ = 0.01). Similar results were found for the contralateral primary motor cortex (M1) hand region of interest (ROI) (see **Appx.A3**).

**Appx. A2.**
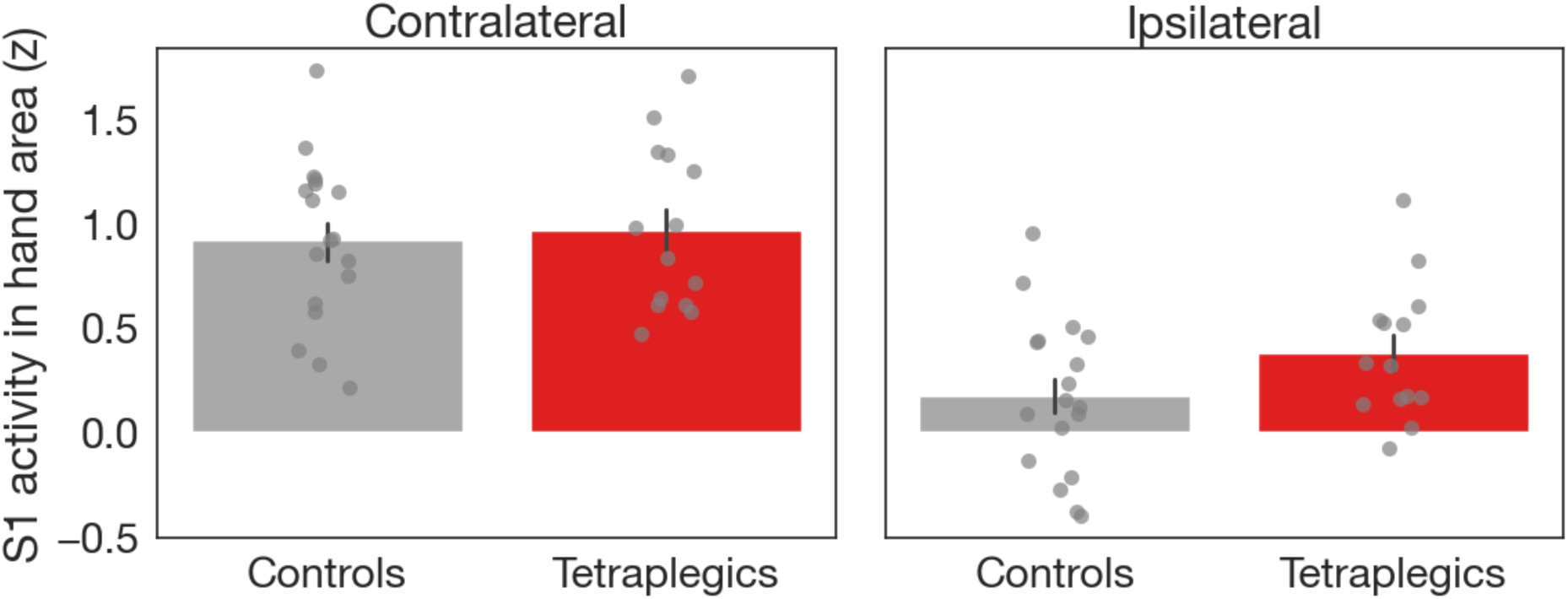
Average movement-related BOLD activity in bilateral S1 of tetraplegic individuals compared to controls. To compare activity levels between hemispheres, activity levels were extracted from both the ipsi- and contralateral S1 hand areas (**Fig. 1**) during attempted unimanual individual finger movements. Tetraplegic individuals significantly activated both their ipsi- and contralateral S1 hand areas, whereas controls predominantly activated their contralateral S1 hand area. Overall task-related activity did not differ significantly between the two groups. As expected, net movement-related activity levels were lower in the ipsilateral than in the contralateral S1 hand area for both groups, and this interhemispheric difference was comparable between controls and tetraplegic individuals. Error bars indicate the standard error of the mean. Dots indicate individual participants.

## APPENDIX A3

**Appx. A3.**
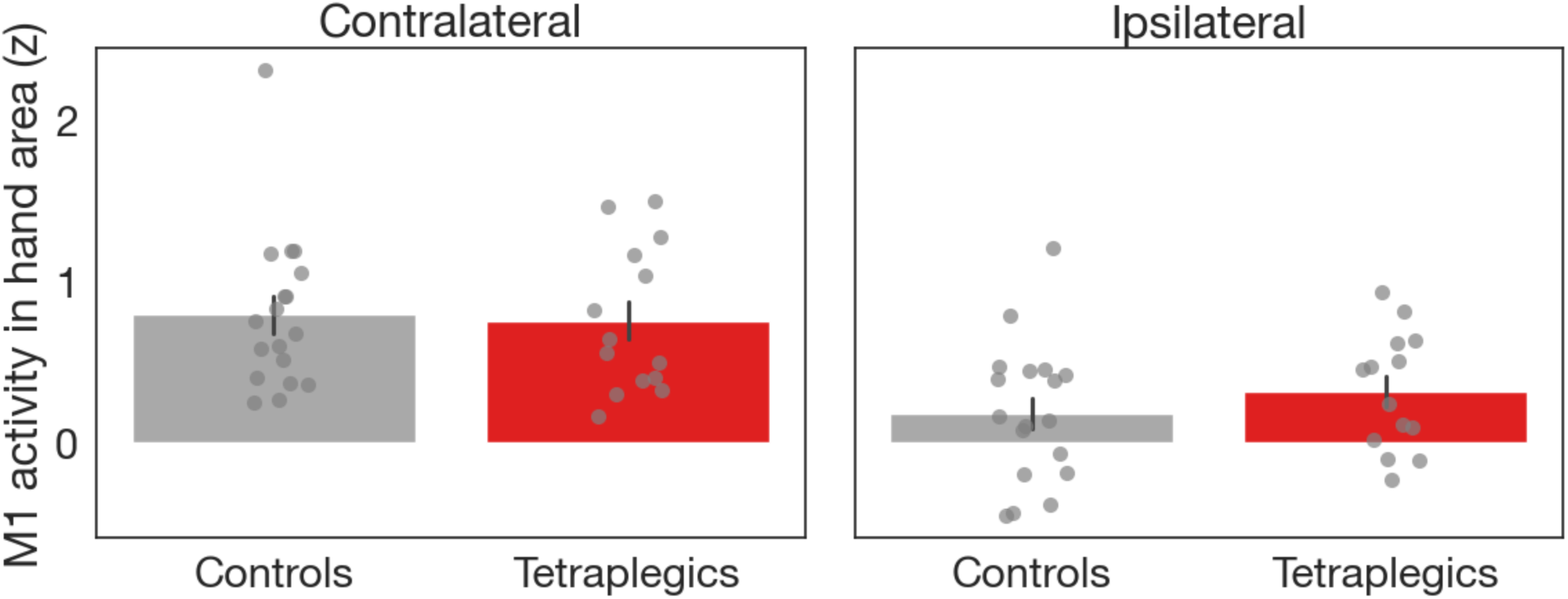
Z-scored BOLD response in bilateral M1 during unimanual movements. Extracted z-scored beta estimates (*ß*) from M1 hand area in ipsilateral and contralateral M1. Again both groups significantly engaged their ipsi- and contralateral M1 hand area by unimanual individual finger movements, except from ipsilateral M1 in controls (controls ipsilateral: t_(17)_ = 1.76, p_corr_ = 0.09, BF_10_ = 0.87; controls contralateral: t_(17)_ = 6.83, p < 0.001, BF_10_ > 100 and tetraplegic individuals ipsilateral: t_(13)_ = 3.23, p < 0.01, BF_10_ = 8.02; tetraplegic individuals contralateral: t_(13)_ = 6.16, p < 0.001, BF_10_ > 100, respectively). A robust mixed ANOVA revealed no significant difference in task-related activity between controls and tetraplegic individuals (F_(1,21.66)_ = 0.09, p = 0.76, η^2^ < 0.05). As expected, beta estimates were lower in the ipsi- than in the contralateral M1 hand area (F_(1,15.38)_ = 148.99, p < 0.001, η^2^ = 0.28). This hemispheric difference in beta estimates trended towards a significant difference between groups (i.e., tendency to interaction effect; F_(1,15.38)_ = 4.00, p = 0.05, η^2^ = 0.01). The black error bars indicate the standard error of the mean. The grey dots represent individual participants.

## APPENDIX A4

**Appx. A4.**
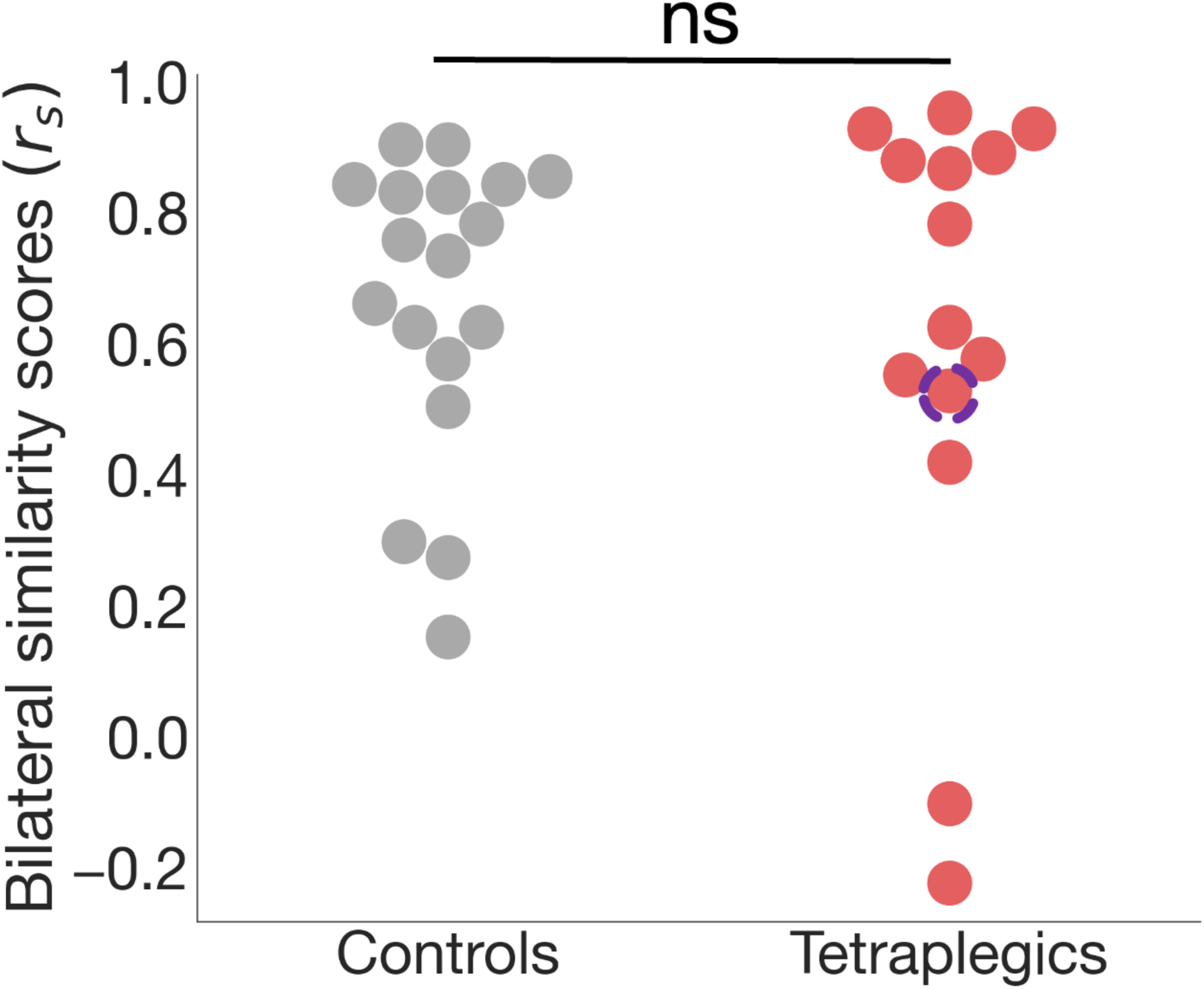
Interhemispheric similarity of finger representational structure in S1. Bilateral similarity (Mantel correlation) between ipsilateral and contralateral S1 hand-area representational distance matrices for controls and individuals with tetraplegia. Each point reflects one participant’s correlation coefficient. There was no significant group difference, although a small number of tetraplegic participants exhibit weak or slightly negative similarity values.

